# Activity of epigenetic inhibitors against *Plasmodium falciparum* asexual and sexual blood stages

**DOI:** 10.1101/694422

**Authors:** Leen Vanheer, Björn F.C. Kafsack

## Abstract

Regulation of gene expression by epigenetic processes is critical for malaria parasite survival in multiple life stages. To evaluate the suitability of targeting these pathways we screened 350 epigenetic inhibitors against asexual blood stages and gametocytes of *P. falciparum*. We observed ≥90% inhibition at 10 µM for 28% of compounds, of which a third retained ≥90% inhibition at 1 µM. These results suggest epigenetic regulation as a promising target for the development of new multi-stage anti-malarials.

---

Despite substantial progress in reducing malaria infections and deaths over the past two decades, the disease remains among the greatest global health challenges, with 219 million cases and 435,000 deaths in 2017 (1). The emergence of drug and insecticide resistance now threatens to reverse these gains and highlights the need for new classes of anti-malarials for use in combination therapies (2). To minimize the emergence and spread of resistance, these new classes should have independent modes of action from existing therapies and be effective against multiple parasite stages, including the asexual blood stages responsible for the disease’s clinical manifestation and gametocytes, the sexual blood stages that mediate transmission.

Recent studies have demonstrated the essential function of multiple genes involved with epigenetic regulation of gene expression in asexual blood stages (3-7). Many of these genes likely also play key roles during the substantial chromatin remodeling that occurs during the early stages of gametocytogenesis (8, 9). Earlier studies involving a limited number of epigenetic inhibitors found activity against malaria parasites (10-14) and several have already been approved for clinical use or are currently in clinical trials for treatment of various cancers (15). To evaluate the promise of targeting epigenetic processes more broadly, we decided to screen the two largest commercially available libraries of epigenetic inhibitors against both asexual blood stages and gametocytes of *Plasmodium falciparum*, the most widespread and virulent human malaria parasite.

Small compound libraries of 209 and 141 epigenetic inhibitors at 10 mM in DMSO were obtained from Selleckchem (Houston, TX) and Cayman chemicals (Ann Arbor, MI), respectively. Libraries were aliquoted and were further diluted in DMSO to 2 mM and 0.2 mM in V-bottom 96-well plates using a Tecan Freedom EVO 150 liquid dispenser (HTSRC, Rockefeller University) and stored at −80°C. A *P. falciparum* NF54 strain expressing the tandem dimeric tomato red fluorescent protein under control of a *peg4* gametocyte promoter (16) was used for cell-based activity screens against both asexual and early gametocytes and maintained at low parasitemia using established methods (17). The activity against asexual blood stages was determined by 72 hour SYBR Green assays (18) as previously described and normalized to solvent-treated controls (included in triplicate on each plate) (19). Activity against developing gametocytes was determined using a flow cytometric assay, as previously described (16). Briefly, highly synchronous asexual cultures were grown for one cycle at 3% haematocrit and 8-9% parasitemia to induce sexual commitment. Percoll-sorbitol isolated schizonts were allowed to invade fresh RBCs for 3-4 hours and remaining late stages were then removed with a second Percoll-sorbitol gradient. The 3-4-hour early rings, both asexual and committed, were seeded into flat-bottom 96-well plates containing compounds at a final 1% haematocrit, 4% parasitemia, 50 mM GlcNAc (Alfa Aesar, Haverhill, MA) in 200 µL and 0.5% DMSO. Cultures were maintained for 6 days before quantifying gametocytemia by flow cytometry (Cytek DxP12). EC50 values were calculated the nlmLS function of the minpack.lm package (v1.2-1) of the R statistical package (v3.6.0).

We screened 350 small molecules known to target epigenetic processes from two commercially available libraries at 10 µM and 1 µM against asexual and early gametocyte blood stages of *P. falciparum*. Seventeen compounds were present more than once, differing only by vendor or counterion. Since responses to repeat compounds showed only minimal variation in response, their mean response is reported (Fig. S1), leaving 332 unique compounds.

Of the compounds screened, 148 had greater than half-maximal activity against at least one stage (Fig. 1 & Fig S3. Also see Fig. S2 and Data Set 1 for activity of all compounds tested). 44% (146) and 16% (54) of compounds had greater than half-maximal activity against asexual stages at 10 µM and 1 µM, respectively (Table 1). Activity against early gametocyte stages was similar, with 36% (120) and 17% (55) of compounds resulting in greater than 50% inhibition at 10 µM and 1 µM, respectively. Greater than 90% inhibition was observed for 28% (92) against asexual blood stages at 10 µM, with 10% (32) retaining ≥90% activity even at 1 µM (Table 2). Against early gametocyte stages 28% (93) and 10% (32) had EC90s below 10 µM and 1 µM, respectively. Despite differences in methodology and parasite strains, these agree well with results for 8 of these compounds that had been screened against either asexual stages or gametocytes in earlier studies (10-14). Thirty-one of the most active compounds (EC90s < 1 µM) were selected for more detailed dose-response studies (Figure 1B). The majority showed similar potency against both blood stages but we found that twelve compounds exhibited at least 2-fold difference in activity against the two parasite stages tested (Fig. 1C). Of the eight compounds more active against asexual stages, seven were histone deacetylase (HDAC) inhibitors while two histone methyltransferase inhibitors, the DNA methyltransferase (DNMT) inhibitor SGI-1027, and the Pan-Jumonji Histone demethylase (HDM) inhibitor JIB-04 (20) were more effective against early gametocytes.

**Figure 1.**
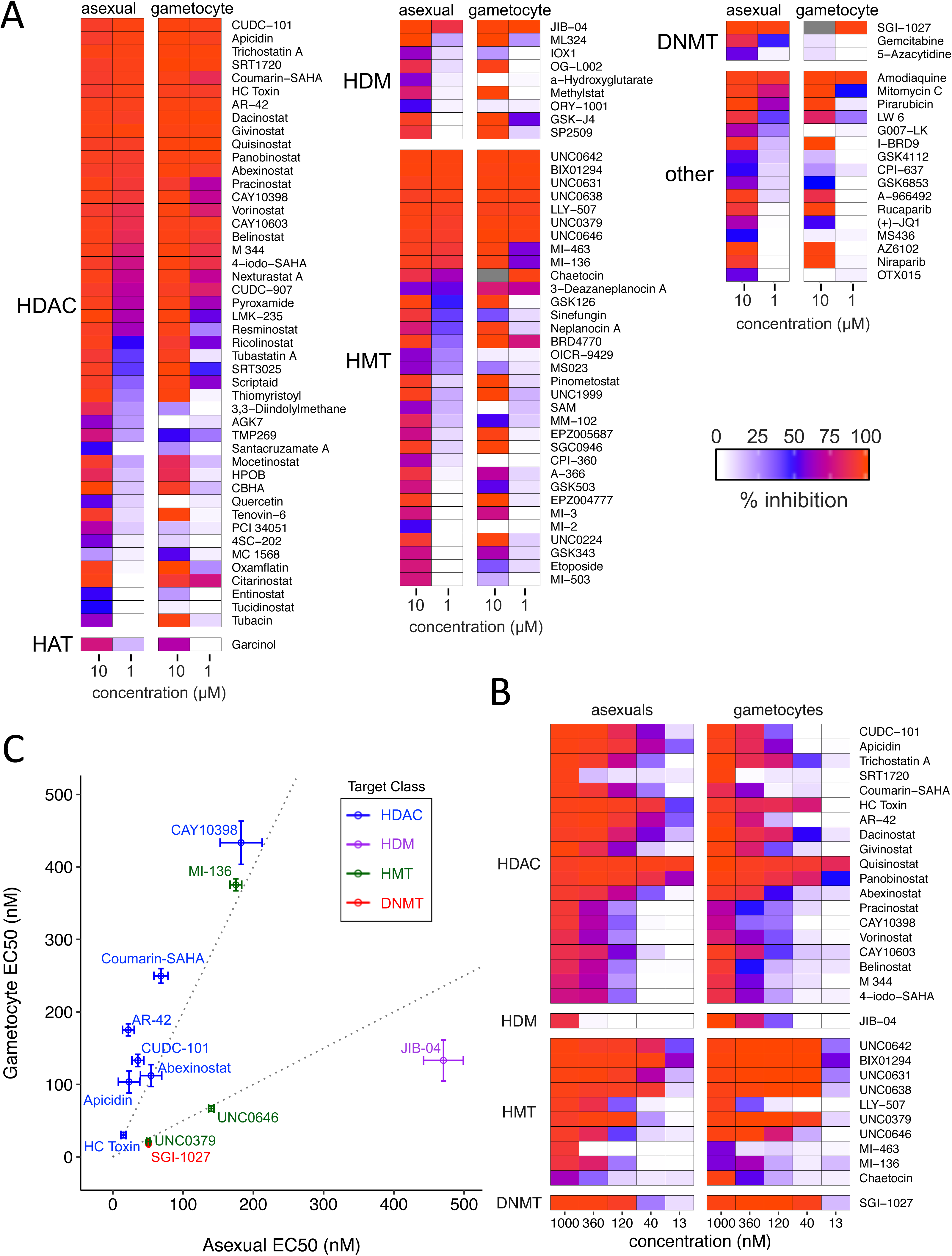
Epigenetic Inhibitors with activity against *P. falciparum* blood stages. **(A)** Compounds with ≥50% inhibition against asexual or early gametocyte blood stages at 10 µM. Heatmap of mean percent inhibition of asexual replication and early gametocyte maturation at 10 and 1 µM compared to solvent-treated controls (n=2, see Table S1 for complete data). Compounds are grouped based on the reported epigenetic process affected in higher eukaryotes: Histone deacetylation (HDAC), histone acetylation (HAT), histone methylation (HMT), Histone Demethylases (HDM), DNA methylation (DNMT), and “other”. Grey color indicates values excluded due to significant hemolysis at 10 µM. **(B)** Additional analysis of dose response for 31 compounds with sub-micromolar EC90s (n=2-3). **(C)** Of these, twelve compounds had a greater than 2-fold difference (indicated by dotted lines) in activity against between asexual blood stages and gametocytes.

**Table 1.** EC50 Activity of epigenetic inhibitors tested grouped by reported epigenetic process targeted in higher eukaryotes.

| Target Class | Number of compounds | >50% inhibition at 10 μM | | >50% inhibition at 1 μM | |
| --- | --- | --- | --- | --- | --- |
|  |  | asexuals | gametocytes | asexuals | gametocytes |
| Histone Acetylation | 10 | 1 (10%) | 1 (10%) | 0 (0%) | 0 (0%) |
| Histone Deacetylation | 89 | 43 (48.3%) | 38 (42.7%) | 25 (28.1%) | 26 (29.2%) |
| Histone Methylation | 52 | 33 (63.5%) | 24 (46.2%) | 11 (21.2%) | 12 (23.1%) |
| Histone Demethylation | 18 | 9 (50%) | 7 (38.9%) | 1 (5.6%) | 2 (11.1%) |
| Histone Phosphorylation | 67 | 41 (61.2%) | 38 (56.7%) | 13 (19.4%) | 13 (19.4%) |
| Histone PARPylation | 24 | 5 (20.8%) | 4 (16.7%) | 0 (0%) | 0 (0%) |
| Histone Reader Domains | 28 | 6 (21.4%) | 3 (10.7%) | 0 (0%) | 0 (0%) |
| DNA Methylation | 15 | 3 (20%) | 1 (6.7%) | 1 (6.7%) | 1 (6.7%) |
| Other | 29 | 5 (17.2%) | 4 (13.8%) | 3 (10.3%) | 1 (3.4%) |
| **Total** | **332** | **146 (44%)** | **120 (36%)** | **54 (16%)** | **55 (17%)** |

**Table 2.** EC90 Activity of epigenetic inhibitors tested grouped by reported epigenetic process targeted in higher eukaryotes.

| Target Class | Number of compounds | >90% inhibition at 10 μM | | >90% inhibition at 1 μM | |
| --- | --- | --- | --- | --- | --- |
|  |  | asexuals | gametocytes | asexuals | gametocytes |
| Histone Acetylation | 10 | 0 (0%) | 0 (0%) | 0 (0%) | 0 (0%) |
| Histone Deacetylation | 89 | 34 (38.2%) | 34 (38.2%) | 15 (16.9%) | 14 (15.7%) |
| Histone Methylation | 52 | 19 (36.5%) | 19 (36.5%) | 9 (17.3%) | 8 (15.4%) |
| Histone Demethylation | 18 | 4 (22.2%) | 6 (33.3%) | 1 (5.6%) | 1 (5.6%) |
| Histone Phosphorylation | 67 | 25 (37.3%) | 26 (38.8%) | 5 (7.5%) | 7 (10.4%) |
| Histone PARPylation | 24 | 4 (16.7%) | 3 (12.5%) | 0 (0%) | 0 (0%) |
| Histone Reader Domains | 28 | 1 (3.6%) | 1 (3.6%) | 0 (0%) | 0 (0%) |
| DNA Methylation | 15 | 1 (6.7%) | 1 (6.7%) | 1 (6.7%) | 1 (6.7%) |
| Other | 29 | 4 (13.8%) | 3 (10.3%) | 1 (3.4%) | 1 (3.4%) |
| **Total** | **332** | **92 (28%)** | **93 (28%)** | **32 (10%)** | **32 (10%)** |

When grouped based on their reported epigenetic targets in higher eukaryotes, compounds that affect the deacetylation, methylation, and phosphorylation of histone in other organisms each had hit rates between 35-40% at 10 µM for both asexual blood stages and gametocytes (Table 1). Genome-wide mutagenesis studies in *P. falciparum* and the rodent malaria parasite *P. berghei* have indicated the essentially of multiple genes encoding histone modifying enzymes (6, 7). While phosphorylation of histone tails has been observed in *P. falciparum* blood stages (21), it remains unclear whether the observed activity of these kinase inhibitors is the result of diminished histone phosphorylation, as the kinases implicated in modification of histone tails in higher eukaryotes also perform other critical functions (see Fig. S3 for kinase inhibitor results).

Hit rates were lower for compounds targeting processes involved in demethylation, acetylation, binding of histone modifications (histone readers), and DNA methylation. *P. falciparum* encodes one or more of genes involved in these pathways and lower hit rates against these may indicate greater divergence from their mammalian homologs or non-essentially of these pathways in blood stages. For example, all but three of the 29 inhibitors of histone readers interfere with the recognition of acetylated histones by Bromo-domains. Earlier studies noted the divergence of these domains in *P. falciparum* while also demonstrating their essentiality for asexual growth. Interestingly, we found that DNA methyltransferase inhibitor SGI-1027 had EC50 values in the low nanomolar range against both stages, despite the fact that the lone DNA methyltransferase in malaria parasites was found to be dispensable for asexual growth in *P. falciparum* (6). When combined with the fact that SGI-1027 contains a quinoline group common in many anti-malarials, this makes an alternative target likely. Overall, our results show that epigenetic regulation of gene expression in malaria parasites is a promising target for interfering with multiple stages of the parasite life cycle.

## Supporting information

Supplemental Data Set 1

## Acknowledgments

We thank the High Throughput and Spectroscopy Resource Center at Rockefeller University for technical assistance, Prof. Photini Sinnis (Johns Hopkins University) for generously providing the NF54 peg4-tdTomato reporter parasites, and Prof. Elisabeth Martinez (UT Southwestern) for additional JIB-04 inhibitor. We also thank L. Kirkman for valuable feedback on the manuscript. This work was supported by a Bohmfalk Charitable Trust Research Grant and NIH R01AI141965 to BK, and a Belgian American Educational Foundation post-doctoral fellowship to LV.

---

**Figure S1.**
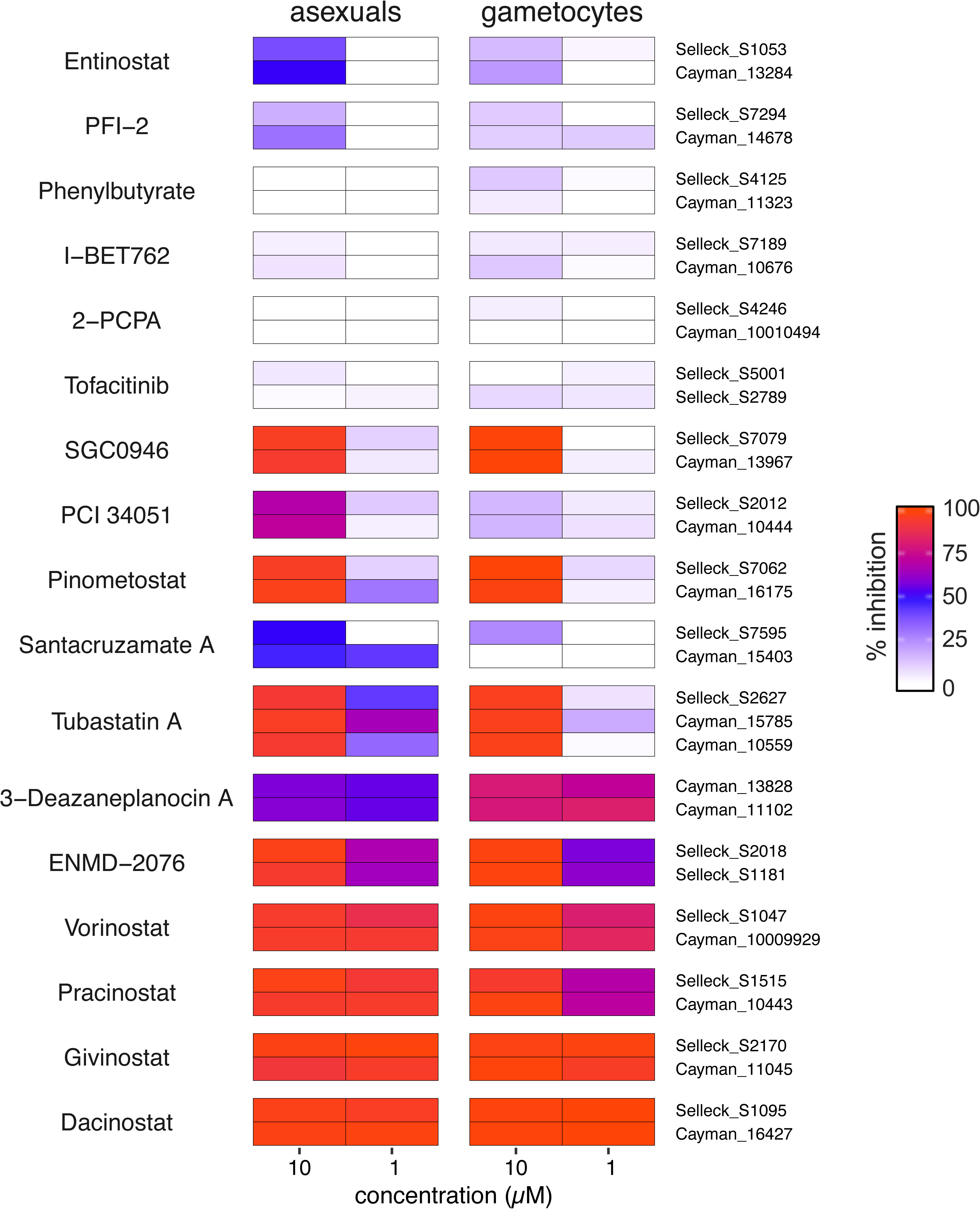
Similar activity of compounds that differed by counterion or vendor.

**Figure S2.**
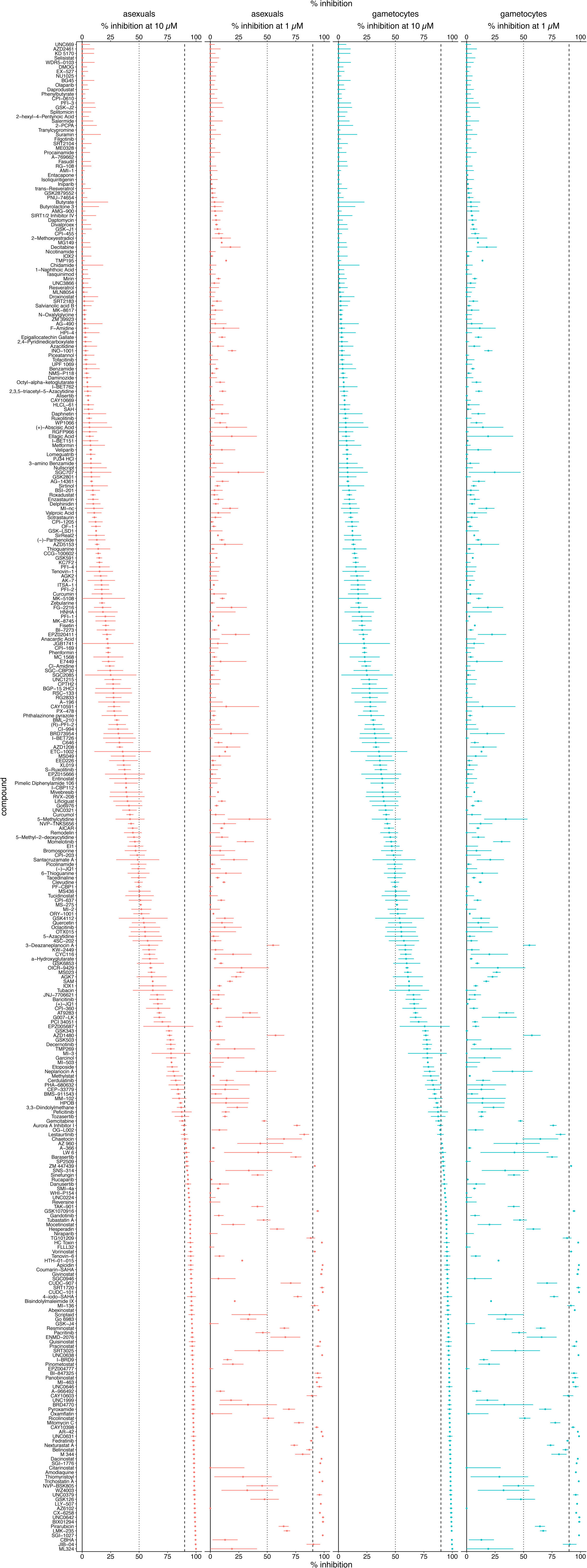
Mean Percent Inhibition of all 337 compounds at 10 µM and 1 µM against asexual blood stages and gametocytes. The dotted and dashed lines indicate 50% and 90% inhibition, respectively. Compounds are ordered by increasing activity against asexual blood stages at 10 µM. Error bars are standard error of n=2.

**Figure S3.**
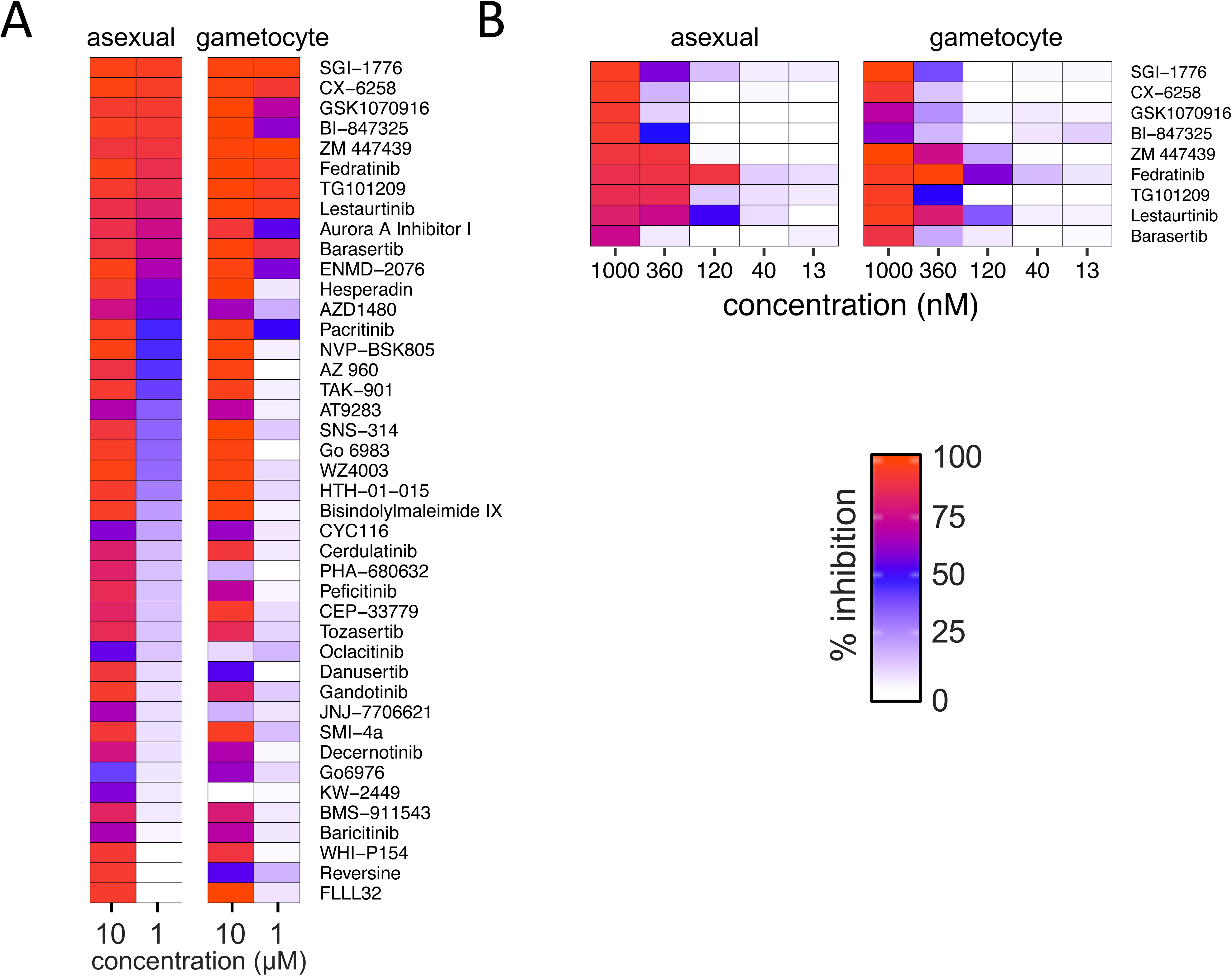
Activity of putative kinase inhibitors. **(A)** Compounds with ≥50% inhibition against asexual or early gametocyte blood stages at 10µM. Heatmap of mean percent inhibition of asexual replication and early gametocyte maturation is shown at 10 and 1 µM compared to solvent-treated controls (n=2, see Table S1 for complete data). **(B)** Additional analysis of dose response for kinase inhibitors with EC90 values below 1 µM (n=2-3).

**Data Set 1.** Mean percent inhibition for all 350 compounds at 10 µM and 1 µM against asexual blood stages and gametocytes, with their observed EC50 range or value.

